# Evaluation of ivermectin effectiveness under contrasting anthelmintic resistance scenarios in beef cattle: a comparative study in Italy and Argentina

**DOI:** 10.64898/2026.07.28.741160

**Authors:** A Bosco, C Canton, M.P. Maurelli, P. Vitiello, L. Ceballos, P. Domínguez, L. Moriones, J. Torres, L. Alvarez, L. Rinaldi, C. Lanusse

## Abstract

Beef cattle farming plays a crucial role in the livestock-related economies of both Italy and Argentina. Although both countries have grazing systems of meat production, the heterogeneity of those systems leads to different parasitological scenarios regarding epidemiology and the presence of resistance to commonly used anthelmintic drugs. Considering ivermectin (IVM) is the most widely used anthelmintic for treating gastrointestinal nematode (GIN) infections in cattle, the current studies evaluated the efficacy of ivermectin (IVM) given subcutaneously (SC) to both young calves and adult cows in ten (10) commercial cattle farms from Italy (A to E) and Argentina (F to J). The work was complemented with the assessment of the IVM plasma exposure and disposition kinetics both in young and adult animals.

While adult cows and young calves were included in the study performed in Italy (Campania region), only calves aged 8-14 months old were selected to perform the efficacy trials in Argentina (central area of the Buenos Aires province). Fifteen (15) calves/cows naturally infected with GINs were treated with IVM (0.2 mg/kg) in each of the Farms. The therapeutic response (efficacy) was determined at 14 days after treatment by the faecal egg count reduction test. Six (6) adult cows and eight (8) young calves treated with IVM were randomly selected to perform the pharmacokinetic (PK) study with blood samples being taken 2 h and 21 days post-treatment. Drug concentrations were measured by HPLC. Similar IVM PK trends were obtained for both adult and young cattle. The IVM systemic exposure (expressed as AUC) obtained for adult cattle (380±158 ng.d/mL) was similar to that observed in calves group (313±85.5 ng.d/mL). No statistical differences between animals from both ages were observed for the most representative PK parameters (P>0.05). Surprisingly, while in Argentina IVM resistance was present in all the farms evaluated (90% CI = 0%-95%), in Italy a fully susceptible nematode population was presented in three farms (90% CI = 96%-100%) and only a very low level of IVM resistance was detected in the rest of the farms (90% CI = 92%-98%). Whereas *Cooperia* spp. and *Haemonchus* spp. were identified as the principal genera exhibiting resistance to IVM in Argentina, *Cooperia* spp., *Ostertagia* spp., and *Oesophagostomum* spp. were reported to display low levels of resistance under the Italian field scenario. In conclusion, IVM can be safely and effectively used in both young and adult cattle with similar disposition kinetic patterns. While its therapeutic response is failing under the conditions investigated in the Argentinian farms, its efficacy remains notably high in the Italian region under study. However, the use of IVM, as well any other active antiparasitic principle, should always be preceded by a diagnostic assessment of the nematode population’s resistance status.

## 1. INTRODUCTION

Beef cattle farming represents a strategic component of livestock production in both Italy and Argentina [1, 2]. However, these countries differ markedly in the type of production systems, environmental conditions, and parasite epidemiology. Argentina is characterized by extensive, pasture-based cow–calf and finishing systems that expose animals to continuous infective larval challenge, thus creating epidemiological conditions highly favorable to the transmission of gastrointestinal nematodes (GINs) [3]. Conversely, Italy presents a more heterogeneous scenario in which extensive grazing systems coexist with semi-intensive and intensive feedlot operations, resulting in variable parasite exposure and infection dynamics across regions [4].

A major challenge shared by both countries is the sustainable control of GIN infections, which significantly affects cattle health, welfare, and productivity. In extensive systems such as those prevalent in Argentina, the environmental contamination with infective larvae leads to high parasite burdens. Under these conditions, the use of anthelmintics, particularly macrocyclic lactones (MLs) such as ivermectin (IVM) has become essential for maintaining animal performance. However, the long-term and often frequent use of IVM in South American grazing systems has accelerated the selection of drug resistant nematode populations, with *Cooperia spp.* and *Haemonchus spp.* representing the most frequently reported resistant genera [5, 6]. Such resistance threatens the efficacy of treatment programs and poses a growing concern for sustainable beef production [7].

In Italy, although epidemiological pressure is generally lower and farming systems more diverse, recent studies and field observations indicate that IVM resistance is emerging and may expand if not addressed through evidence-based control strategies [8, 9]. This situation highlights the need for harmonized pharmacological and parasitological assessments capable of supporting informed decision-making in helminth control.

IVM remains the most widely used broad-spectrum anthelmintic to control GIN infection in cattle. As a ML compound, IVM exhibits potent activity against adult and immature stages of most GINs, including hypobiotic larvae, lungworms, and several ectoparasites [10, 11]. Its efficacy is strongly influenced by pharmacokinetic (PK) processes, which determine drug concentrations at the parasite–host interface and ultimately shape treatment outcomes [12, 13, 14]. In this context, plasma concentration–time profiles following subcutaneous administration represent a reliable predictor of the drug’s availability in tissues where nematodes reside.

Given the importance of aligning drug exposure with parasite susceptibility and considering the contrasting production systems and resistance scenarios between Italy and Argentina, integrated pharmaco-parasitological evaluations are essential. Recent comparative studies have emphasized the need to characterize both the PK behavior and field efficacy of IVM across diverse cattle categories and farming contexts [15]. Such knowledge is fundamental to developing sustainable parasite control programs that preserve drug efficacy, limit resistance development, and optimize cattle health and productivity.

The current studies aimed to perform an integrated pharmaco-parasitological evaluation of IVM in beef cattle raised under contrasting production systems in Italy and Argentina were part of a bi-national collaborative research project. The specific objectives were to (i) assess the field efficacy of subcutaneous IVM against GINs in naturally infected cattle from commercial farms in both countries, using standardized faecal egg count reduction tests (FECRT) to identify the presence and extent of anthelmintic resistance; (ii) characterize the plasma PK profiles of IVM following subcutaneous administration in different cattle categories (young animals vs. adult cows) to determine if physiological and management-related factors may influence drug disposition and systemic exposure; (iii) compare GIN resistance patterns between Italian and Argentine farms, considering their different features and production systems.

## 2. MATERIALS AND METHODS

### 2.1 Parasitological trial

#### 2.1.1 Field trial and animals

Field trials were conducted on a total of ten commercial beef cattle farms located in Italy and Argentina. Specifically, in Italy, five beef cattle farms (A - E) were included and selected in the Campania region (southern Italy); cattle were crossbreeds (Limousine, Podolica, Marchigiana) cows and calves aged 6-12 months and 1-3 years old, respectively. In Argentina, five farms (F - J) were included and selected in the Humid Pampean Region; cattle were Aberdeen Angus calves aged 8– 14 months old. Both in Italy and Argentina, beef production was based on a grazing system, and all the selected animals were naturally infected with GINs.

The selection of cattle farms was mainly driven by the willingness of the farmers and by the representativeness of the area concerning GIN infection levels. All farms included in this trial had a history of ivermectin use in previous years.

On day −1 on each farm, 20-25 animals were checked for GIN egg per gram (EPG) counts, ear-tagged, and the individual body weights were recorded. Different EPG inclusion thresholds were applied (Italy: ≥20 EPG; Argentina: ≥100 EPG), reflecting the distinct epidemiological conditions and baseline infection levels in the two countries. Experimental animals had an average of 53 and 782 EPG counts in Italy and Argentina, respectively. All the animals had free access to water. Animal procedures and management protocols were approved by the Ethics Committee (act 11/2020) of the Facultad de Cs. Veterinarias, Universidad Nacional del Centro de la Provincia de Buenos Aires (UNCPBA), Tandil, Argentina. In Italy, the study was conducted in accordance with the guidelines of the Declaration of Helsinki and the Ethics Committee of the University of Naples. Informed consent was obtained from all farm owners involved in the study.

#### 2.1.2 Coprocultures

The coproculture analysis was performed following the protocol of the UK Ministry of Agriculture, Fisheries and Food [16] to assess the composition of GIN genera before and after treatment. For each group of cattle, pooled faecal samples were prepared at Day 0 and Day 14 by combining equal amounts of faeces from individual animals. The pooled samples were incubated at room temperature (23-26°C) for 14 days to allow the development of third-stage larvae (L3).

Subsequently, L3 were recovered and morphologically identified using the keys described by van Wyk and Mayhew [17]. The identification and relative abundance of each nematode genus were determined by examining 100 L3 per sample; when fewer than 100 larvae were recovered, all available larvae were identified. This approach allowed the estimation of the proportional representation of each genus within the total larval population of each group.

#### 2.1.3 Faecal Egg Count Reduction Test

The anthelmintic efficacy of the different treatments was assessed by the FECRT, according to the recommendations of the last World Association for the Advancement of Veterinary Parasitology (WAAVP) guidelines [18]. On each farm, 9-20 animals were treated with IVM injectable solution (Ivomec®, 1% solution, Boehringer Ingelheim Animal Health, Italy or Argentina) at the dosage of 0.2 mg/kg. Faecal samples were individually collected directly from the rectum of each cow/calf during pre-treatment (day −1) and again on day 14 post-treatment to determine faecal egg count reduction (FECR) post-treatment [18]. Mini-FLOTAC technique with a detection limit of 5 EPG [19] was used to analyze the faecal samples and estimate EPG counts. The data analysis was conducted using the FECRT web-based platform (www.fecrt.com), applying the delta method as described by [20].

In addition, efficacy against different genera was calculated by partitioning the mean faecal egg count of each treatment group pre and post treatment, by the proportion of L3 of each genus in the corresponding coproculture [21].

### 2.2 PK trial

#### 2.2.1 Field trial and animals

The PK trial was carried out in Argentina, both in cows and calves. Six cows and eight calves treated with IVM (the same formulation and dose used in the FECRT) were used in the PK trial. Blood samples (10 mL) were taken from the jugular vein in heparinized Vacutainer tubes (Becton Dickinson, NJ, USA) as follows: before treatment and at 2, 4, 6, 8 h and 1, 2, 3, 7, 15 and 21 days post-treatment. Plasma was separated by centrifugation at 3000 g for 15 min, placed into plastic tubes and frozen at −20°C until analysis by High Performance Liquid Chromatography (HPLC).

#### 2.2.2 IVM analytical procedures

The extraction of IVM from spiked and experimental plasma samples was carried out following an adaptation of the technique described by [22]. An aliquot of 0.25 mL of plasma sample was combined with doramectin (DRM) (used as internal standard) and then 1 mL of acetonitrile was added to each sample. After mixing for 20 min, samples were sonicated in an ultrasonic bath for 10 min (Transonic 570/H, Laboratory Line Instruments Inc., Melrose Park, IL, USA). The solvent-sample mixture was centrifuged at 2000 g during 15 min and the supernatant was manually transferred into a tube and concentrated to dryness under a stream of nitrogen. The resuspension was carried out with a solution of N-methylimidazole (Sigma Chemical, St. Louis, MO, USA) in acetonitrile (1:1) [23]. Derivatization was initiated by adding trifluoroacetic anhydride (Sigma Chemical, St Louis, MO, USA) solution in acetonitrile (1:2). Finally, an aliquot of this solution was injected directly into the chromatographic system. IVM concentrations were determined by HPLC using a Shimadzu 10 A-HPLC system with a fluorescence detector (Shimadzu, RF-10 Spectrofluorometric detector, Kyoto, Japan). Calibration curves were prepared in the range between 0.2 and 100 ng/mL. The linear regression lines for IVM showed correlation coefficients ≥0.99. The LOQ was established at 0.2 ng/mL. The mean recovery percentages for concentrations ranging between 0.2 and 100 ng/mL (n = 6) was 93% with CV of 8.5%.

#### 2.2.3 Pharmacokinetic and statistical analysis of the data

Data concentration profiles for IVM obtained after the treatment of each individual animal were analyzed using a non-compartmental approach with version 2.0 of the PkSolutions software (Summit Research Service, CO, US). The peak concentration (Cmax) and time to peak concentration (Tmax) were recorded directly from the measured concentration data. Pharmacokinetic parameters were determined. The elimination half-life (T½el) and absorption half-life (T½ab) were calculated as ln2/ß and ln2/k, respectively, where ß represent the terminal slope (h^−1^) and k is the slope obtained by feathering which represents the first order absorption rate constant. The rates were calculated by performing regression analysis using data points belonging to the terminal or absorption phase concentration-time plot. The area under the plasma concentration-time curve from zero up to the quantification limit (AUC_0-t_) was calculated using the trapezoidal rule [24] and further extrapolated to infinity (AUC_0-∞_) by dividing the last experimental concentration by the terminal elimination rate constant (ß). Statistical moment theory was applied to calculate the mean residence time (MRT) according to Perrier and Mayersohn [25].

The PK parameters and concentration data are reported as arithmetic mean ± Standard Deviation (SD). PK parameters for IVM calculated after its administration to cows and calves were statistically compared using Student t-test. A value of P<0.05 was considered statistically significant. The statistical analysis was performed using the Instat 3.0 software (Graph Pad Software, CA, USA).

## 3. RESULTS

### 3.1 Parasitological trial

Table 1 presents the overall faecal egg counts recorded across all farms at day 14 post-treatment, along with the lower and upper 90% confidence intervals and the corresponding nematode population status. Firstly, EPG counts observed in both countries showed a higher level of nematode infection in Argentina (286-1452 EPG) compared to Italy (37-67 EPG). Regarding nematode population status, on Italian farms A, D, and E, the 90% confidence intervals ranged between 95.8% and 100%, indicating that the nematode populations on these farms were susceptible to IVM. Farms B and C showed a low level of resistance, with 90% confidence intervals ranging from 91.9% to 98.5%. In contrast, on all Argentine Farms including in the study the analysis of the 90% CI indicated that the animals were infected with IVM-resistant nematodes. In fact, the lower 90% confidence interval ranged from −22% to 83% across all farms.

**Table 1.**
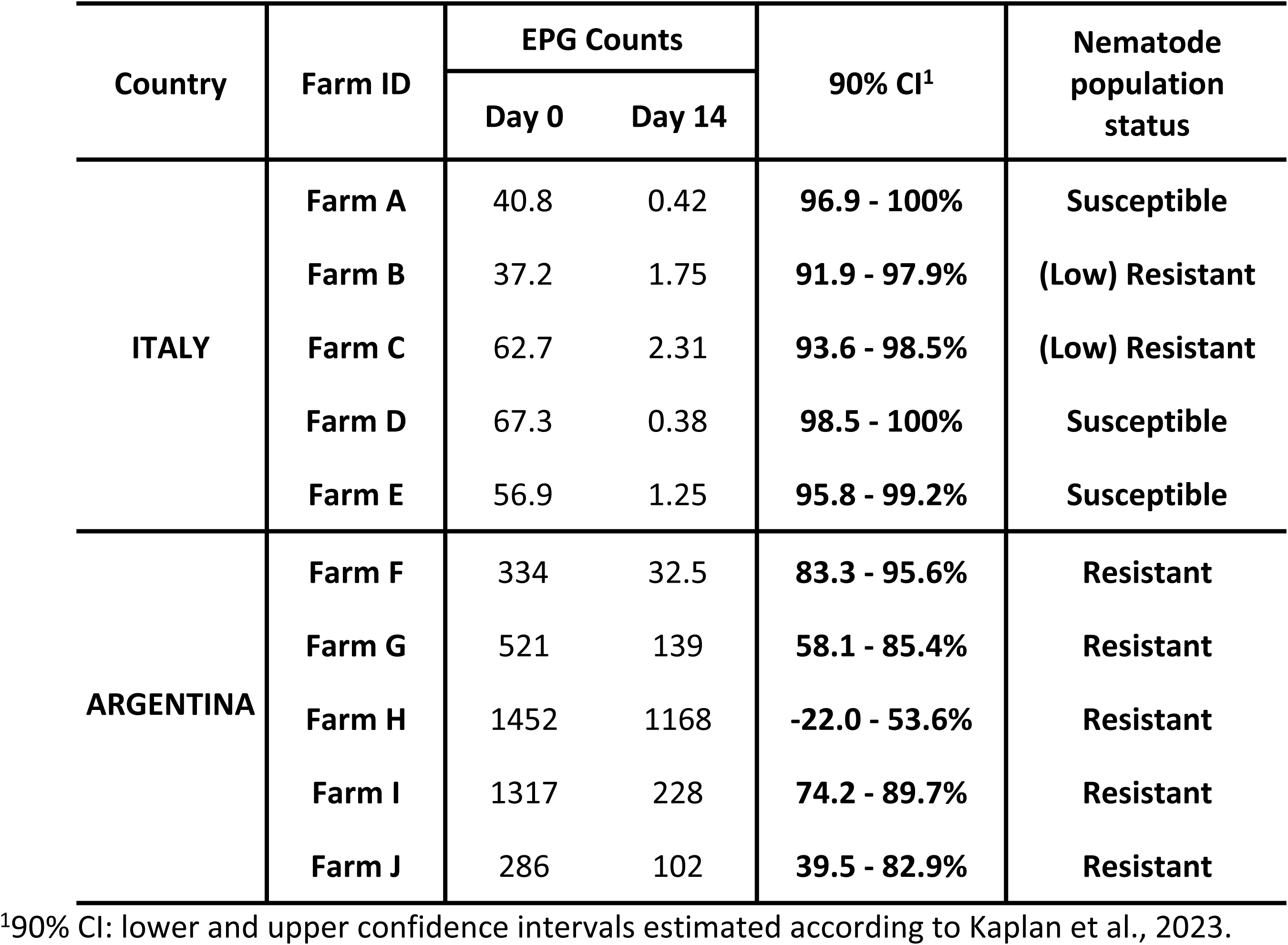
Nematode egg per gram counts (EPG, arithmetic mean), lower and upper confidence intervals 90%, and nematode population status after the subcutaneous (SC) administration of ivermectin (IVM, 0.2 mg/kg) to naturally parasitized cattle in Italy and Argentina.

The anthelmintic efficacies against *Cooperia* spp., *Haemonchus* spp., *Ostertagia* spp., *Oesophagostomum* spp. and *Trichostrongylus* spp. in both countries are shown in Table 2. On Italian farms in which the overall efficacy indicated a susceptible scenario, IVM treatment effectively controlled all GIN genera. In contrast, on farms B and C, where the analysis of overall efficacy indicated a low level of resistance, IVM failed to adequately control *Cooperia* spp., *Ostertagia* spp. and *Oesophagostomum* spp., with efficacies ranging between 93.4% and 94.5%. In Argentina, *Cooperia* spp. and *Haemonchus* spp. were the main genera showing resistance to IVM. Notably, IVM failed to control *Cooperia* spp. on all farms included in the study, with markedly lower efficacies than those observed in Italy, ranging from 56% to 82%. Additionally, and in contrast to the Italian farms, IVM-resistant *Haemonchus* spp. were detected on four Argentine farms, with efficacies ranging from 0% to 92%. In contrast, subcutaneous administration of IVM on Argentine farms achieved effective control of *Ostertagia* spp., *Oesophagostomum* spp., and *Trichostrongylus* spp. across all farms included in the study.

**Table 2.** Reduction percentages of faecal egg counts (FECRT) for *Cooperia, Haemonchus*, *Ostertagia*, *Oesophagostomum* and *Trichostongylus* spp. (based on egg counts partitioned to genera using the proportion of each genus recovered as larvae from faecal larval cultures) after the subcutaneous (SC) administration of ivermectin (IVM, 0.2 mg/kg) to naturally parasitized cattle in Italy and Argentina

| Genus | FECRT Day 14<sup>1</sup> |  |  |  |  |  |  |  |  |  |
| --- | --- | --- | --- | --- | --- | --- | --- | --- | --- | --- |
|  | ITALY |  |  |  |  | ARGENTINA |  |  |  |  |
|  | Farm A | Farm B | Farm C | Farm D | Farm E | Farm F | Farm G | Farm H | Farm I | Farm J |
| *Cooperia* spp. | 100% | 93.4% | 94.1% | 100% | 98.4% | 81.2% | 77.7% | 56.8% | 68.9% | 82.0% |
| *Haemonchus* spp. | 100% | - | - | 100% | - | 100% | 67.8% | 0% | 67.5% | 92.0% |
| *Ostertagia* spp. | 100% | 97.5% | 93.7% | 100% | 97.9% | 96.0% | 100% | 100% | 100% | 97.0% |
| *Oesophagost.* spp. | 100% | 94.5% | 93.9% | 100% | 96.8% | 100% | 100% | 100% | 100% | 100% |
| *Trichostrong.* spp. | 100% | - | 98.4% | 100% | 98.4% | 100% | - | - | - | - |

<sup>1</sup>FECR estimated according to McKenna (1990).

### 3.2 PK Trial

Figure 1 presents the mean (± SD) plasma concentration profiles for IVM after the SC administration to calves and cows. IVM plasma levels were measured up to 21 days post-treatment. Similar IVM PK trends were obtained for both adult and young cattle. Table 3 summarizes the plasma PK parameters obtained for cows and calves for IVM. No statistical differences between both categories were observed for the most representative PK parameters (P>0.05). In fact, the IVM systemic exposure (expressed as AUC_0-t_) obtained for adult cows (380±158 ng.d/mL) was similar to that observed in young calves group (313±85.5 ng.d/mL).

**Figure 1.**
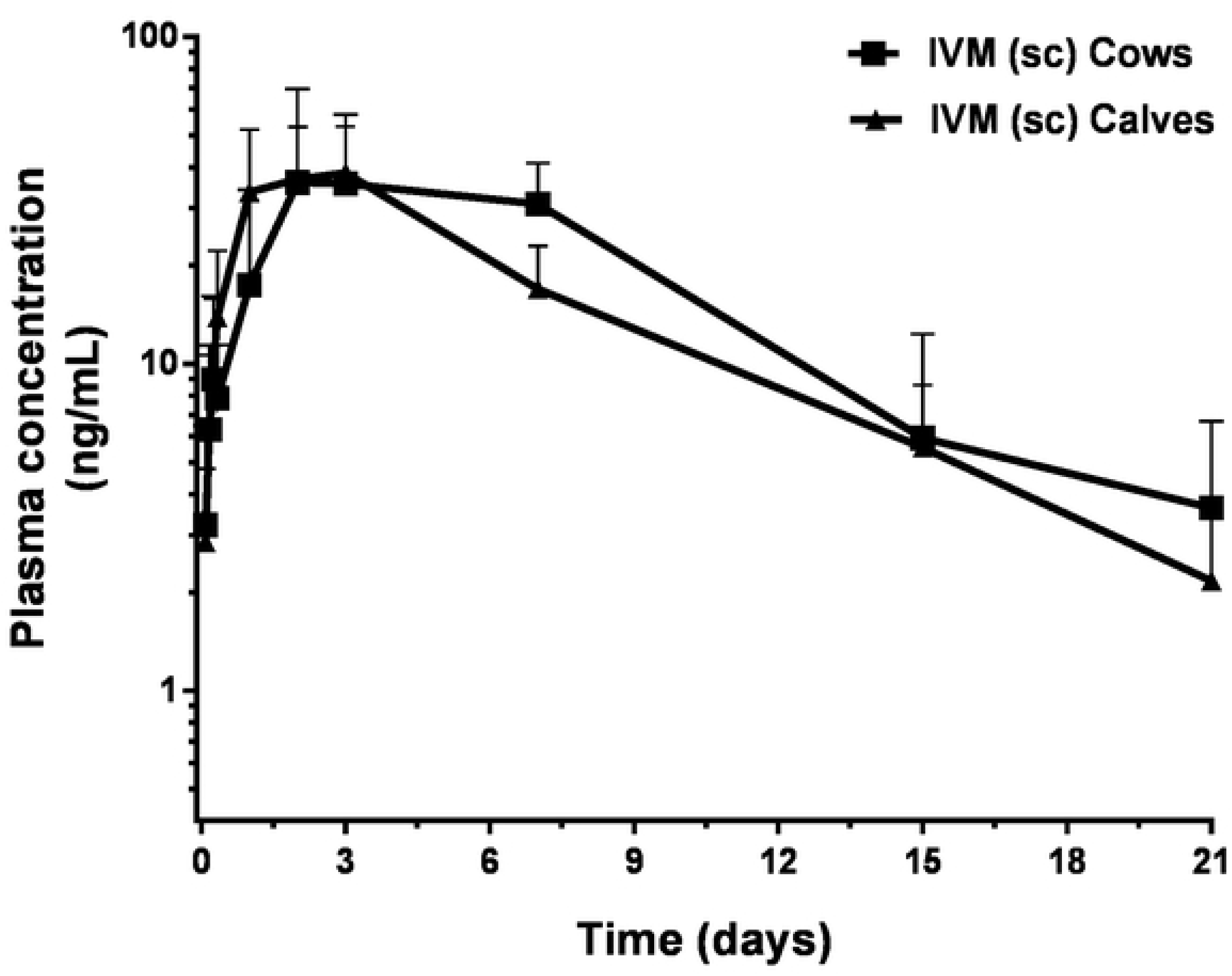
Nematode egg per gram concentration profiles obtained after its subcutaneous administration (0.2 mg/kg) to parasitized cows (n=6) or calves (n = 8).

**Table 3.** Plasma pharmacokinetic parameters (mean ± SD) for ivermectin (IVM) obtained after its subcutaneous administration (0.2 mg/kg) to beef cows and calves.

| IVERMECTIN |  |  |
| --- | --- | --- |
| Pharmacokinetic parameters | COWS | CALVES |
| Tmax (d) | 4.20 ± 2.59<sup>a</sup> | 2.63 ± 0.74<sup>a</sup> |
| Cmax (ng/mL) | 42.4 ± 29.6<sup>a</sup> | 41.9 ± 17.5<sup>a</sup> |
| AUC<sub>0-t</sub>(ng·d/mL) | 380 ± 158<sup>a</sup> | 313 ± 85.5<sup>a</sup> |
| AUC<sub>0-∞</sub>(ng·d/mL) | 412 ± 159<sup>a</sup> | 329 ± 86.7<sup>a</sup> |
| MRT (d) | 8.46 ± 3.45<sup>a</sup> | 6.96 ± 1.99<sup>a</sup> |
| T<sub>1/2</sub>ab (d) | 1.42 ± 0.48<sup>a</sup> | 0.72 ± 0.42<sup>b</sup> |
| T<sub>1/2</sub>el (d) | 4.65 ± 2.29<sup>a</sup> | 4.48 ± 1.52<sup>a</sup> |

IVM: 0.2 mg/kg BW (sc). Tmax: time to peak plasma concentration; Cmax: peak plasma concentration; AUC<sub>0-t</sub>: area under the concentration vs. time curve from time zero to the quantification time; AUC<sub>0-∞</sub>: area under the concentration-time curve extrapolated to infinity; MRT: mean residence time; T<sub>1/2</sub>ab: absorption half-life; T<sub>1/2</sub>el: elimination half-life. Pharmacokinetic parameters with different superscript letters are statistically different (P<0.05).

## 4. DISCUSSION

IVM is the most widely used anthelmintic to treat GINs in cattle worldwide. The studies reported here provide a comparative pharmaco-parasitological evaluation of IVM in beef cattle raised under contrasting production systems in Italy and Argentina. By simultaneously assessing the plasma disposition and systemic exposure of subcutaneously administered IVM, and its field efficacy against GINs, these studies offer an integrated perspective essential for developing sustainable parasite control strategies.

Consistent with previous research, IVM demonstrated high efficacy in most Italian farms, reflecting a production system characterized by lower grazing pressure and more heterogeneous management practices. The reduced environmental larval availability and the less intensive use of MLs in Italy likely contribute to a slower selection for resistant nematode populations. These findings align with recent European surveys reporting limited but emerging IVM resistance in cattle, particularly in *Cooperia spp.* [26, 27, 8]. This phenomenon has been widely reported over the last decade in major cattle-rearing countries in Europe, including Italy, Germany, France, and the UK [28]. In that study, decreased efficacy of MLs was observed in a high number of farms in all countries except Italy (IVM, n = 24; moxidectin, n = 20). In France, reduced efficacy of both IVM and moxidectin was mainly associated with the dose-limiting nematode species *Cooperia* spp., whereas in the UK and Germany resistance involved not only dose-limiting species but also the more pathogenic abomasal nematode *Ostertagia ostertagi*. Species-specific efficacy calculations in the European study confirmed that reduced efficacy predominantly affected *Cooperia* spp.; however, in approximately 25% of farms, efficacy against *O. ostertagi* was also below 95%. In agreement with these findings, the present study identified low-level resistance primarily associated with *Cooperia* spp. on two Italian farms (93–94% efficacy), while reduced efficacy against *Ostertagia* spp. (93% efficacy) was detected only on one farm. Overall, although IVM resistance appears to be less prevalent, the pattern observed in Italy mirrors the broader European scenario, with *Cooperia* spp. remaining the main driver of reduced macrocyclic lactone efficacy and only sporadic involvement of *Ostertagia* spp.

In contrast, several farms in Argentina exhibited reduced IVM efficacy, with *Cooperia spp.* and *Haemonchus spp.* confirmed as the predominant resistant genera—an observation fully consistent with previous reports from South American grazing systems [29, 5, 30]. The extensive, pasture-based environments typical of Argentina promote near-constant exposure to infective nematode larvae, resulting in higher parasite burdens and frequent anthelmintic treatments. This ecological and management context has long been recognized as a driver of accelerated resistance selection, particularly to MLs such as IVM [31]. The elevated frequency of anthelmintic treatments administered during the first grazing season has likely played a pivotal role in the selection and establishment of resistant parasite populations in Argentina. Under these conditions, repeated whole-group treatments are often applied in the absence of significant parasite challenge, markedly reducing the refugia population. This management scenario creates strong selection pressure, favoring the survival and subsequent predominance of resistant genotypes within the nematode population. In Argentina, resistance to IVM was detected in 93% of the farms included in a nationwide survey performed several years back [5]. Consistent with the findings of the present study, the principal genera associated with IVM resistance were *Cooperia* spp. and *Haemonchus* spp. Notably, both investigations reported the presence of IVM-resistant *Cooperia* spp. in 100% of the farms where IVM resistance was identified. Additionally, IVM failed to control *Haemonchus* spp. in 4 of the 5 farms included in the study. An IVM failure to control these GINs is consistent with previous reports in Argentina [6, 32] and in Brazil [29, 33]. Since *Cooperia* spp. is a dose limiting species for IVM, this is the genus in which IVM resistance would be first expected [34]. Notably, the negative impact of inadequate control of resistant GINs on cattle productivity is well established, both in infections involving IVM-resistant *Cooperia* spp. alone [35] and in mixed infections with IVM-resistant *Cooperia* spp. and *Haemonchus* spp. [7, 32].

From a pharmacokinetic standpoint, plasma concentration–time profiles in both calves and adult cows were consistent with the well know behavior of subcutaneously administered IVM, characterized by prolonged absorption, extensive distribution between plasma and tissues and low metabolism rate, as is typical for MLs. The time of parasite exposure to active drug concentrations determines the efficacy and/or persistence of activity for their activity in ruminants [14]. Mean plasma concentration-time profiles obtained in the current study were similar to those reported for IVM in previous trials in cattle [12, 6]. As previously described, subcutaneously administered IVM is slowly absorbed, attaining peak plasma concentrations (Cmax) of 42.4 ng/mL in cows and 41.9 ng/mL in calves at 2–4 days post-treatment, and demonstrating extensive systemic exposure, as reflected by a mean residence time of approximately 7-8 days (see Table 1). Importantly, the IVM systemic exposure (expressed as AUC_0-t_) obtained for adult cows (380±158 ng.d/mL) was similar to that observed in young calves group (313±85.5 ng.d/mL). The only significant difference between the two bovine categories was observed in the absorption half-life (T_½_ab), reflecting a faster absorption in calves (0.72 d) compared to cows (1.42 d). In contrast, no effect of animal age/category was observed on the elimination half-life (T_½_el). These results are consistent with those reported by Toutain et al. [36] Importantly, this slight difference in absorption rate is unlikely to have any clinical impact on IVM efficacy in either cows or calves. In fact, no statistical differences between both categories were observed for the most representative PK parameters (P>0.05). Therefore, the PK profiles were comparable between young animals and adult cows, confirming that age-related differences in body weight and metabolic activity did not substantially alter systemic drug exposure under field conditions.

Instead, the combined interpretation of parasitological and pharmacological data strongly supports the conclusion that treatment failures arise primarily from true drug resistance rather than insufficient drug exposure. This finding underscores the necessity of coupling PK assessment with field efficacy testing when investigating suspected resistance, as PK alterations and resistance can lead to similar clinical outcomes but require fundamentally different management responses.

The cross-country comparison highlights the value of a harmonized methodological approach. Assessing efficacy and PK profiles simultaneously across diverse production systems allows for a more robust interpretation of drug performance and provides actionable insights for refining parasite control programs. In particular, the high efficacy observed in Italy suggests that sustainable practices - including periodic monitoring of resistance, reduced treatment frequency, and evidence-based interventions-can maintain IVM effectiveness over time. Conversely, the scenario in Argentina illustrates the consequences of long-term, frequent anthelmintic use and emphasizes the urgency of implementing diagnostic-guided treatment protocols.

Overall, IVM can be safely and effectively used at recommended doses in both young cattle and adult cows. While *Cooperia* spp. and *Haemonchus* spp. were the main genera resistant to IVM in most of the Argentinian farms, its efficacy remains notably high at the assessed Italian farms. The combined pharmaco-parasitological framework adopted in these studies demonstrates that IVM remains an effective anthelmintic when susceptibility is preserved, but its benefits cannot be guaranteed without regular diagnostic assessment of the nematode population’s resistance status. The sustained efficacy observed across Italian farms reinforces the importance of continued diagnostic monitoring to prevent the spread of resistance, especially in regions where intensification or repeated treatments may enhance selection pressure. Ensuring the sustainable use of IVM requires the integration of surveillance tools, responsible treatment practices, and tailored management strategies that account for the specific epidemiological context of each production system. The scientific evidence generated through this bi-national collaboration underscores the need for continued efforts to develop complementary and interdisciplinary research in animal health and related fields.

## ACKNOWLEDGEMENTS

This study was funded by the bi-national collaborative research project “Prog. Coop. Bilateral Nivel 1 CONICET-CUIA 2019” (Project number 24120190100127CO) from Argentina and Italy. The authors would like to thank the farmers for collaborating with this study.

